# Genome mining as a biotechnological tool for the discovery of novel biosynthetic genes in lichens

**DOI:** 10.1101/2022.05.04.490581

**Authors:** Garima Singh, Francesco Dal Grande, Imke Schmitt

**Author notes:** **Corresponding author:** Garima Singh. **Emails:** Garima Singh, Francesco Dal Grande, Imke Schmitt.

## Abstract

The ever-increasing demand for novel drugs highlights the need for bioprospecting unexplored taxa for their biosynthetic potential. Lichen-forming fungi (LFF) are a rich source of natural products but their implementation in pharmaceutical industry is limited, mostly because the genes corresponding to a majority of their natural products is unknown. Furthermore, it is not known to what extent these genes encode structurally novel molecules. Advance in next-generation sequencing technologies has expanded the range of organisms that could be exploited for their biosynthetic potential. In this study, we mine the genomes of nine lichen-forming fungal species of the genus *Umbilicaria* for biosynthetic genes, and categorize the BGCs as “associated product structurally known”, and “associated product putatively novel”. We found that about 25-30% of the biosynthetic genes are divergent when compared to the global database of BGCs comprising of 1,200,000 characterized biosynthetic genes from planta, bacteria and fungi. Out of 217 total BGCs, 43 were only distantly related to known BGCs, suggesting they encode structurally and functionally unknown natural products. Clusters encoding the putatively novel metabolic diversity comprise PKSs (30), NRPSs (12) and terpenes (1). Our study emphasizes the utility of genomic data in bioprospecting microorganisms for their biosynthetic potential and in advancing the industrial application of unexplored taxa. We highlight the untapped structural metabolic diversity encoded in the lichenized fungal genomes. To the best of our knowledge, this is the first investigation identifying genes coding for NPs with potentially novel therapeutic properties in LFF.

## Background

Natural products (NPs) are small molecules in nature produced by the organism. Historically, NPs have played a key role in drug discovery due to their broad pharmacological effects encompassing antimicrobial, antitumor, anti-inflammatory properties and against cardiovascular diseases [1,2]. In the past decades about 70% of the drugs were based on NPs or NP analogs [1,2]. The demand for novel drugs however, is ever increasing due to the emergence of antibiotic-resistant pathogens, the rise of new diseases, the existence of diseases for which no efficient treatments are available yet, and the need for replacement of drugs due to toxicity or high side-effects [3,4]. One way to address global health threats and to accelerate NP-based drug discovery efforts is bioprospecting unexplored taxa to assess their biosynthetic potential and identify potentially novel drug leads.

Genes involved in the synthesis of a NPs are often grouped together in biosynthetic gene clusters [5–7]. These clusters have a core gene which codes for the backbone structure of the NP and other genes which may be involved in the modification of the backbone or may have a regulatory or transport-related function [5,8–10]. Depending upon the core gene, the BGCs could be grouped into the following major classes: non-ribosomal peptide synthetases (NRPS), polyketide synthases (PKS), NRPS-PKS (hybrid non-ribosomal peptide synthetase-polyketide synthase), terpenes, and RiPP (ribosomally synthesized and post-translationally modified peptide). Conserved motives, especially of the PKS genes, facilitate the bioinformatic detection of the clusters [11–14].

Traditionally, a large portion of NP-based drugs have been contributed by a few organisms as the drug discovery was mostly restricted to culturable organisms [15–17]. In the last decades, bioinformatic prediction of biosynthetic gene or biosynthetic gene clusters (group of two or more genes that are clustered together and are involved in the production of a secondary metabolite) has revolutionized NP-based drug discovery as this process is culture-independent and enables rapid identification of entire biosynthetic landscape from so far unexplored NP resources, including silent or unexpressed genes. Two tools have been vital to bioinformatic approach to drug discovery: AntiSMASH [18] and MIBiG [19]. AntiSMASH includes one of the largest BGC database for BGC prediction [18] whereas MIBiG (Minimum Information about a Biosynthetic Gene Cluster) is a data repository allowing functional interpretation of target BGCs by comparison with BGCs with known functions [19]. Recently, efforts have been made to cluster homologous BGCs into gene cluster families (GCFs) and to simultaneously identify novel BGCs [20,21]. Two tools have been introduced to cluster BGCs into GCFs: BiG-FAM clusters structurally and functionally related BGCs into GCFs and identifies structurally most diverse BGCs by comparing the query BGCs to about 1,200,000 BGCs of the BiG-FAM database [21]. BiG-SLiCE clusters homologous BGCs of a dataset into GCFs without reference to an external database, to identify unique BGCs in it [20]. Bioinformatic prediction and clustering of BGCs allows rapid identification of potentially novel drug leads, reducing the costs and time associated with drug discovery by early elimination of unlikely candidates.

Lichens, symbiotic organisms composed of fungal and photosynthetic partners (green algae or cyanobacteria, or both), are suggested to be treasure chests of biosynthetic genes and NPs [22–24]. Although the number of identified NPs per LFF is typically less than 5 [25], the number of BGCs in the genomes of LFF may range from 25-60 [12]. It is not well known how BGCs from LFF relate in structure and function to BGCs from bacteria and non-lichenized fungi, i.e., if a portion of the BGC landscape of LFF is distinct, and might serve as a source of NPs with novel therapeutic properties. Difficulties associated with heterologous expression of LFF genes have so far restricted the application of LFF-derived NPs in the industry. Recently two biosynthetic genes from LFF have been successfully heterologously expressed [9,26]. This, combined with advances in long-read sequencing technology (higher genome quality), and low cost of sequencing provide a promising way forward to discover LFF-derived NPs with pharmacological potential.

Here we employ a long-read sequencing based comparative genomics and genome mining approach to estimate the BGC functional diversity of nine species of the lichenized fungal genus *Umbilicaria*. Specifically, we aim to answer the following questions: (1) What is the functional diversity of BGCs in *Umbilicaria*? and 2) what is the percentage of novel BGCs and species-specific BGCs in *Umbilicaria*?

## Results

### Overview of BGCs in the *Umbilicaria* genomes

*Umbilicaria* genomes contain 20-33 BGCs, with the highest number of BGCs detected in *U. deusta* and lowest in *U. phaea* (Fig. 1A). We did not observe a correlation between genome size and number of BGCs (correlation coefficient = 0.10). *Umbilicaria* species contain an average of 13 PKS clusters, and 4.2 NRPS clusters per species (Fig. 1B), making a PKS to NRPS clusters proportion of 3.1). The most dominant class of BGC in *Umbilicaria* are the ones with PKSs, amounting more than 50% of the total BGCs, followed by terpene clusters (about 20%) and NRPS clusters (about 15%) respectively, (Fig. 2A). In contrast, NRPSs are the most dominant class among fungal and bacterial BGCs (Fig. 2B, C), amounting to about 42% and 30% respectively.

**Fig 1.**
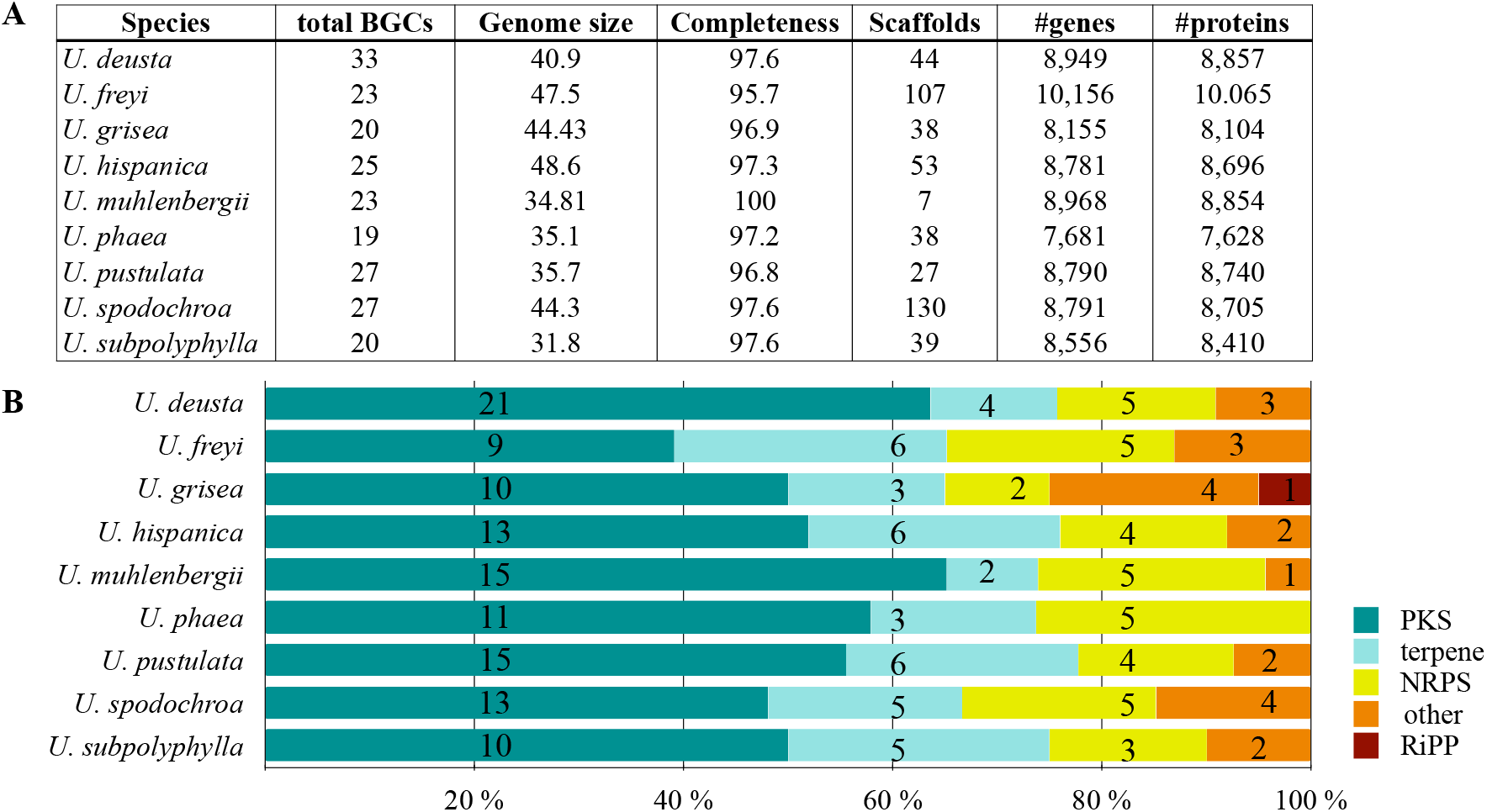
Genome quality metrics and diversity of biosynthetic genes in nine species of *Umbilicaria*. **A)** Genome metrics including the total number of biosynthetic gene clusters as predicted by antiSMASH, and number of genes and proteins estimated by InterProScan and SignalP as implemented in the funannotate pipeline. **B)** Diversity of biosynthetic gene clusters associated with major natural product categories, indicated as percentages (colored bars) and absolute numbers (numbers on bars).

**Fig 2.**
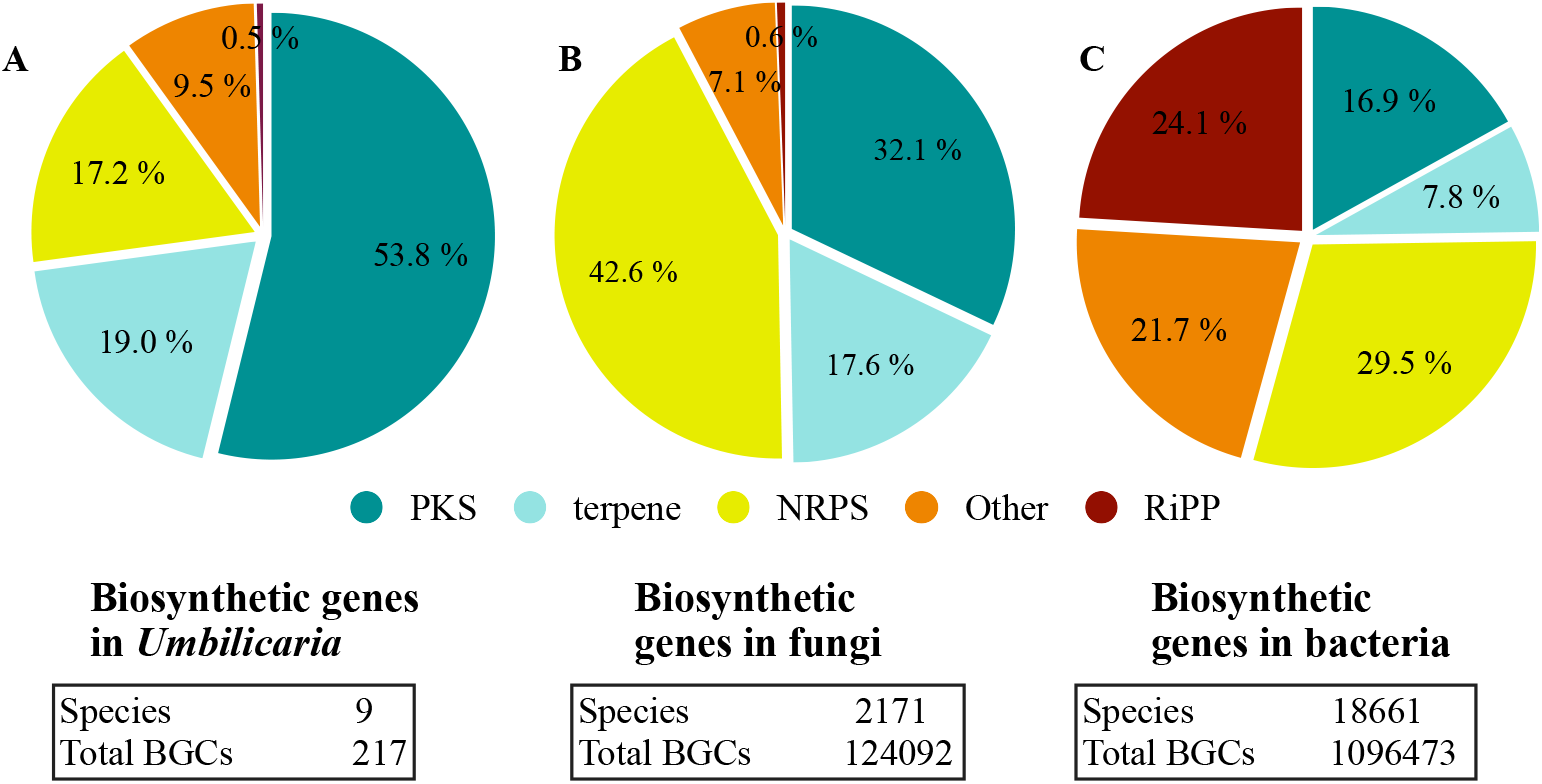
Biosynthetic gene clusters in **A)** *Umbilicaria*, **B)** the full fungal BGC dataset and **C)** full bacterial BGC dataset. PKSs are the most dominant class of BGCs in *Umbilicaria* whereas in fungi and bacteria NRPSs are the predominant BGC class. Although the publicly available LFF genomes (> 50) are much lower than the non-lichenized fungi (about 2100), all the LFF genomes analyzed for their BGCs have PKSs as the most common class of BGCs (see discussion for details), suggesting that the predominance of PKSs as observed here in *Umbilicaria* dataset is a common feature of LFF genomes.

### BGC clustering: BiG-FAM

Of the total 217 BGCs found in 9 *Umbilicaria* species, 18 BGCs (8%) obtained a BGC-to-GCFs (Gene Cluster Families) pairing distance lower than 400, indicating that they potentially code for structurally very similar compounds known from the BGCs of their respective GCFs (Fig. 3A, B); 156 (71%) had a pairing distance of 400-900, suggesting that they share similar domain architectures with previously described BGCs in the BiG-FAM database. We identify the clusters belonging to above two groups as “associated product structurally known”. 43 BGCs (21%) had a pairing distance greater than 900 and are potentially BGCs encoding novel natural products (Fig. 3 A). We identify these clusters as “associated product putatively novel”. These BGCs belong to the class terpenes (1 BGCs), NRPSs (12 BGCs) and PKSs (30 BGCs). The details of these BGCs and the sequence of the core gene is provided in the Additional file 1.

**Fig. 3.**
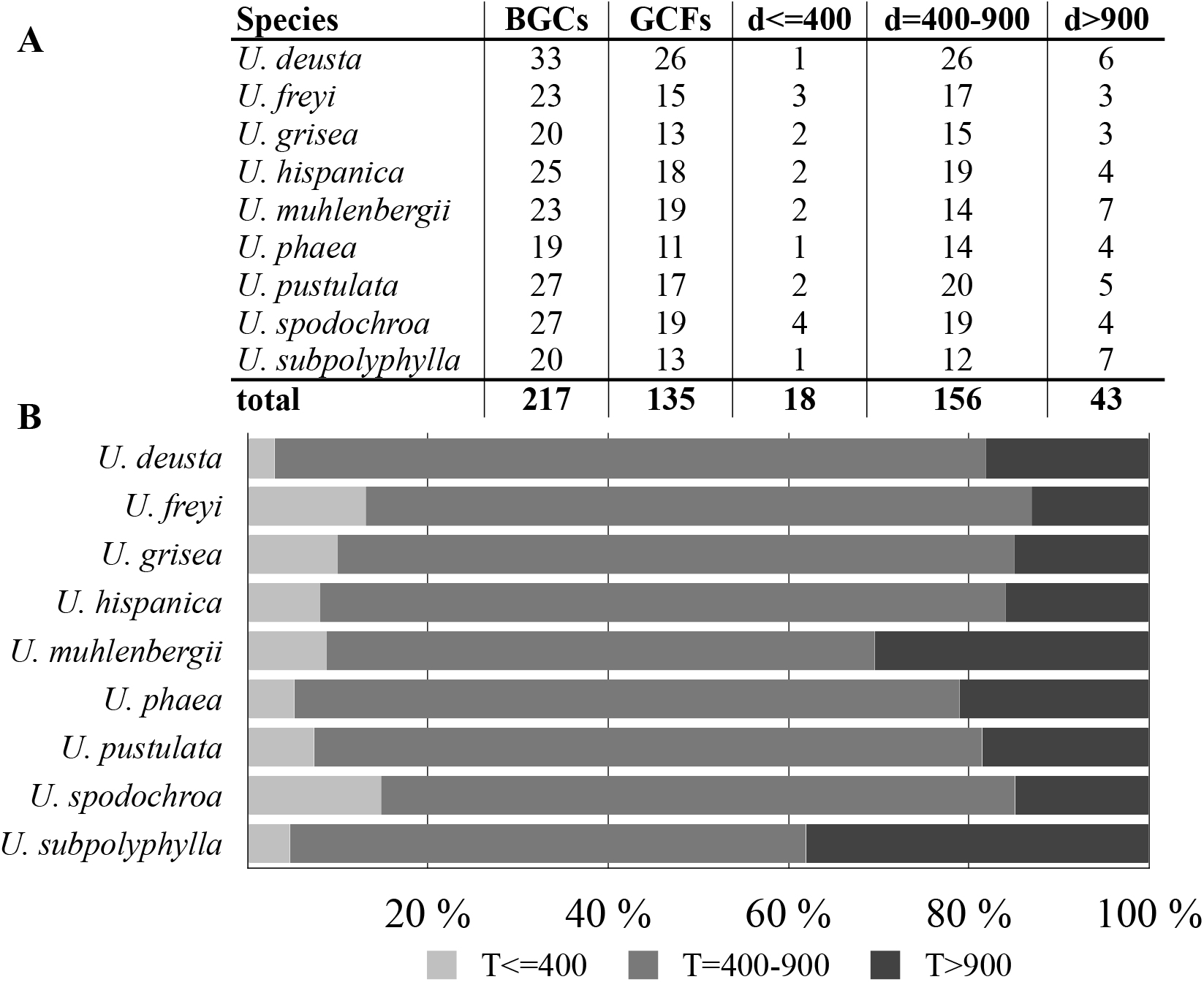
**A)** Total BGCs in *Umbilicaria* and GCFs as identified by BiG-FAM and the number of BGCs clustering into a pre-characterized gene cluster families (GCFs) in BiG-FAM and their distance groups. d<=400 suggest that the cluster codes for a structurally and functionally similar NP, d=400-900 indicates that the BGC codes for a related but structurally and functionally divergent NP, whereas d>900 suggests that the BGC codes for a novel NP. **B)** Bar plots representing the percentage of BGCs in each *Umbilicaria* species with d<= 400, d= 400-900 and d>900. Only a small proportion of BGCs in each species could be grouped into a pre-characterized GCF in the BiG-FAM database (21,678 species, 1,225,071 BGCs and 29,955 GCFs), whereas a large proportion of them is only distantly related to the pre-characterized BGCs. About 15-30% of BGCs could not be grouped into BiG-FAM gene cluster families and potentially code of structurally and functionally divergent NPs.

### Within-genus comparison of BGCs: BiG-SLiCE

We identified species-specific BGCs within *Umbilicaria* using BiG-SLiCE. Out of 217 total BGCs, 159 (72%) grouped into 20 GCFs (d=900), suggesting they are similar clusters shared by multiple species, while 58 (28%) had a d > 900, indicating that they were only distantly related to other BGCs in *Umbilicaria*. Each *Umbilicaria* species contains four to ten (6.45 – 16.13%) unique, species-specific BGCs (Additional file 2A). In *U. deusta* we detected two BGCs (both with PKSs) that were extremely divergent (d > 1800) within the genus (Additional file 2B).

Out of these BGCs, 15 are unique within *Umbilicaria* as well divergent from the BGCs to the known BGCs present in BiG-FAM database.

## Discussion

Lichens produce a large number of natural products, and they have even more BGCs [27–29]. However, whether these BGCs encode hitherto unknown metabolic diversity/chemical structures is not known. Here we quantify, for the first time, the proportion of BGCs linked to putatively novel natural products in a group of closely related lichen-forming fungi. The identification of 23 clusters encoding putatively novel chemical structures can be useful in the search for new structures and drug leads.

In this study we mined the genomes of the *Umbilicaria* spp. to identify all the BGCs (Fig. 1), followed by clustering the structurally and functionally similar BGCs into gene cluster families (Fig. 3A, B) and identifying the gene clusters potentially coding for novel NPs (Fig. 4, Additional File 1). Using *Umbilicaria* spp. as a study system, we show that LFF biosynthetic landscape is diverse from that of non-lichenized fungi and bacteria, being particularly rich in PKSs (Fig 2) and that a substantial portion for LFF BGCs (about 28% in case of *Umbilicaria*) potentially codes for novel NPs (Fig. 3A, B). To the best of our knowledge, this is the first investigation of this kind, implementing state of the art computational tools to determines the proportion of metabolic diversity in LFF coding for novel drugs and identifying candidate genes as a source of drug leads to prioritize them for drug discovery efforts.

**Fig. 4.**
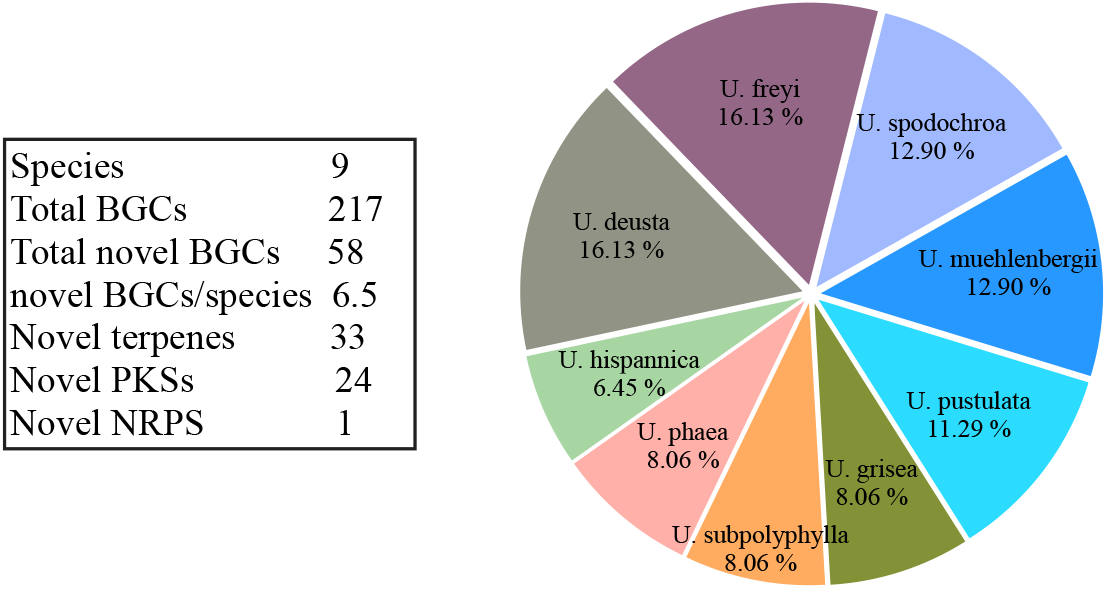
Pie chart depicting the contribution of each species to the overall novel *Umbilicaria* BGCs (as identified by BiG-SLiCE, T>900) Each *Umbilicaria* species contains about 4-10 unique, species-specific BGCs. *U. freyi* and *U. deusta* contain the highest number of novel BGCs. The number of novel BGCs slightly positively correlated to the number of clusters (R=0.68). Out of 58 BGCs unique BGCs (T>900) 56.89% were terpene- and 41.37% were PKS clusters.

### Biosynthetic potential and BGC diversity of *Umbilicaria* spp

Although only PKSs-derived NPs are reported from *Umbilicaria* species (gyrophoric-, umbilicaric-, and hiascic acid etc.) [30–32], we found that the *Umbilicaria* BGC landscape is biosynthetically diverse and comprises three to five classes of NPs (Fig 1A, B). This is also the case for most other LFF, for instance, PKS-derived NPs, are reported from *Bacidia* spp., *Cladonia* spp., *Endocarpon* spp., *Evernia prunastri, Umbilicaria pustulata, Pseudevernia furfuracea*, but all of them contain several PKS, NRPS and terpene gene clusters [12,29,32–34]. All these above-stated studies show that the biosynthetic potential of LFF vastly exceeds their detectable chemical diversity. On average LFF may contain up to 30-40 BGCs but the number of identified compounds per species is usually less than 10 [12,33,35]. This could be because most of the clusters are silent and do not synthesize the NP or it could be simply because of the failure to detect the NP. Bioinformatic characterization of entire BGC landscape followed by identification of most distinct BGCs provides a way to estimate the novelty of all the BGCs including the unexpressed and silent ones.

### BGC diversity of LFF as compared to bacteria and non-lichenized fungi

We identified five classes of BGCs in the *Umbilicaria* genomes. PKSs were the most dominant class, amounting to about 50%, followed by terpenes (19%), and NRPSs (14%) (Fig. 1, Fig. 2 A). BGCs including PKSs typically make up the majority of BGCs in LFF: *Evernia prunastri* (60%), *Pseudevernia furfuracea* (61%), *Cladonia* spp. (65%), *Endocarpon pusillum* (58%), *Lobaria pulmonaria* (46%), and *Ramalina peruviana* (63%) (cite).

Although the number of publicly accessible, good quality LFF genomes are rather scarce for LFF (<25) as compared to the bacteria and non-lichenized fungi, the data available (9 *Umbilicaria* spp. genomes [36] plus 9 other publicly available lichen genomes) suggests that the predominance of PKSs is a common feature of BGCs in LFF contributing more than 50% to the total BGCs. In contrast, in bacteria and non-lichenized fungi, NRPS are the most prevalent BGC class, amounting to about 30% and 42% respectively, followed by the PKSs (Fig. 2 B, C). This suggests that the biosynthetic potential of LFF is unique as compared to the other organisms traditionally exploited for NPs, i.e., non-lichenized fungi and bacteria, especially with respect to PKS diversity. In this regard, a recent study suggested that although bacteria and fungi may share a few NPs, they do not have an overlapping chemical space and instead have distinct biosynthetic potential [37]. LFF having a distinct BGC landscape presents a complementary resource of NPs with promising medicinally-relevant biosynthetic properties.

### *Umbilicaria* BGCs: Gene Cluster Families (GCFs) and novel NPs

Gene cluster families (GCFs) are the groups of BGCs that encode the same or very similar molecules. A total of 217 BGCs from nine *Umbilicaria* species were clustered into of 135 unique GCFs. (Fig 3 A) This suggests that *Umbilicaria* spp. are potentially capable of synthesizing many structurally and functionally different natural products, although in nature only one compound class is typically detected (depsides, linked to a BGC containing a PKS).

Only a small fraction of *Umbilicaria* BGCs, 8%, could be clustered with the pre-characterized BGCs (Fig. 3A, B). About 71% of the BGCs were clustered to the BiG-FAM GCFs with d= 400-900, indicating that they were only distantly related in structure and function (Fig. 3 A, B). These BGCs are also interesting candidates to be investigated for their biosynthetic properties as even a minor difference in the cluster and the chemistry of the final metabolites could cause a crucial difference in bioactivity related to function and the pharmacological potential of the product [38].

About 21% percent BGCs were highly divergent (d>900) and are novel, potentially coding for structurally and functionally unique NPs and could be an interesting target for NP-based drug discovery (Fig. 3 B). The strikingly high number of novel BGCs in a fungal genus adds to the mounting evidence that the non-model and understudied taxa are enormous, untapped source of novel NPs.

Genome mining for large genomic regions, such as fungal BGCs, works best when the genomes under study are highly complete and contiguous, as well as reliably annotated. Many publicly available LFF genomes do not fulfill these criteria, preventing a taxonomically broad study of biosynthetic novelty encoded in the genomes of LFF. We were surprised that even a “chemically boring” lichen taxon, such as the genus *Umbilicaria*, harbored 43 BGCs encoding putatively unknown natural product diversity. It lets us suspect that chemically more diverse taxa, e.g. Lecanorales or Pertusariales, each including hundreds of species, are even richer sources of BGCs with novel functions, and compounds with potential pharmaceutical applications. Increased genome sequencing of taxonomically diverse LFF, combined with higher genome qualities will facilitate BGC discovery.

### Unique BGCs within *Umbilicaria spp*.: BiG-SLiCE

BGCs which are uniquely occurring in a species are candidates for interesting NPs [20,37,39]. On average each *Umbilicaria* species contains seven species-specific BGCs. Most of the novel BGCs are present in *U. deusta* and *U. freyi* whereas *U. hispanica* has lowest number of novel BGCs (Fig. 4). This suggests that even closely related species (species within a single genus) contain diverse biosynthetic potential. Species or strain specific biosynthetic potential has already been demonstrated for LFF, for example in *Umbilicaria pustulata* [32] and *Pseudevernia furfuracea* [33] and it is a rather common occurrence in fungi [32,37,40]. For instance, majority of the BGCs in *Streptomyces*, i.e., 57% have been shown to be strain-specific [41]. The unique BGCs within *Umbilicaria* belong to the BGC classes PKSs, terpenes, NRPS as well as to indoles (Supplementary information S2). Of these, mostly only PKS derived NPs have been well studies in LFF and shown to have diverse pharmacological properties [42–44].

Two PKS obtained a pairing distance greater than 1800. These were the most divergent BGCs (Supplementary information S2) within *Umbilicaria* and were “orphan BGCs”, i.e., for these clusters the corresponding metabolite cannot be predicted. Recently several orphan clusters have been activated to synthesize a compound with useful pharmacological properties, for example the antibiotic holomycin gene cluster from the marine bacterium *Photobacterium galatheae* was activated in culture [45–48]. The novel and orphan clusters reported in this study are potentially interesting candidates for synthesizing molecules with unique pharmacological properties and may serve as drugs leads.

About 17% of fungal BGCs, 8% of bacterial BGCs and 19% of LFF BGCs comprise terpenes (Fig. 2). Terpenes are pharmaceutically extremely versatile, with antimicrobial, anti-inflammatory, neurodegenerative, and cytotoxic properties [49–54]. Some of the common plant-derived terpenes and terpenoids are curcumin, Eucalyptus oil. Although several studies reported pharmacological properties of fungal terpenes, such studies on LFF are missing despite the slightly larger proportion of terpenes in LFF genomes. In this study we report structurally and functionally unique terpenes as promising candidates, to be investigated for their pharmaceutical potential.

## Conclusion

In this study we identified the biosynthetic diversity of the lichen forming fungal genus *Umbilicaria*, grouped the structurally and functionally related clusters into GCFs and identified the most diverse, potentially novel clusters. Using *Umbilicaria* as model system we show that LFF constitute a valuable source of novel NPs suggesting that there is tremendous natural product diversity to be discovered in them. In particular they are rich source of novel PKSs and terpenes. Combining this observation with other sequenced LFF we show that LFF are indeed a source of untapped natural product diversity.

## Materials and methods

### Dataset

The genomes of the following *Umbilicaria* species were used for this study: *U. deusta, U. freyi, U. grisea, U. subpolyphylla, U. hispanica, U. phaea, U. pustulata, U. muhlenbergii* and *U. spodochroa*. Except *U. muhlenbergii* which belongs to the Bioproject PRJNA239196, all the other genomes are a part of Bioproject PRJNA820300 (Table 1). The details of sample and library preparation, as well as genome sequencing for *U. muhlenbergii* are available in Park et al. [55] and for the other eight *Umbilicaria* spp in Singh et al. [36]. Briefly, all the genomes except *U. muhlenbergii* were generated via PacBio SMRT sequencing on the Sequel System II using the continuous long read (CLR) mode or the circular consensus sequencing (CCS) mode. The continuous long reads (i.e. CLR reads) were then processed into highly accurate consensus sequences (i.e. HiFi reads) and assembled into contigs using the assembler metaFlye v2.7 [56]. The contigs were then scaffolded with LRScaf v1.1.12 (github.com/shingocat/lrscaf, [57]). We used only binned Ascomyocta reads for this study (extracted using blastx in DIAMOND (--more-sensitive --frameshift 15 –range-culling) on a custom database and following the MEGAN6 Community Edition pipeline [58]).

**Table 1.**
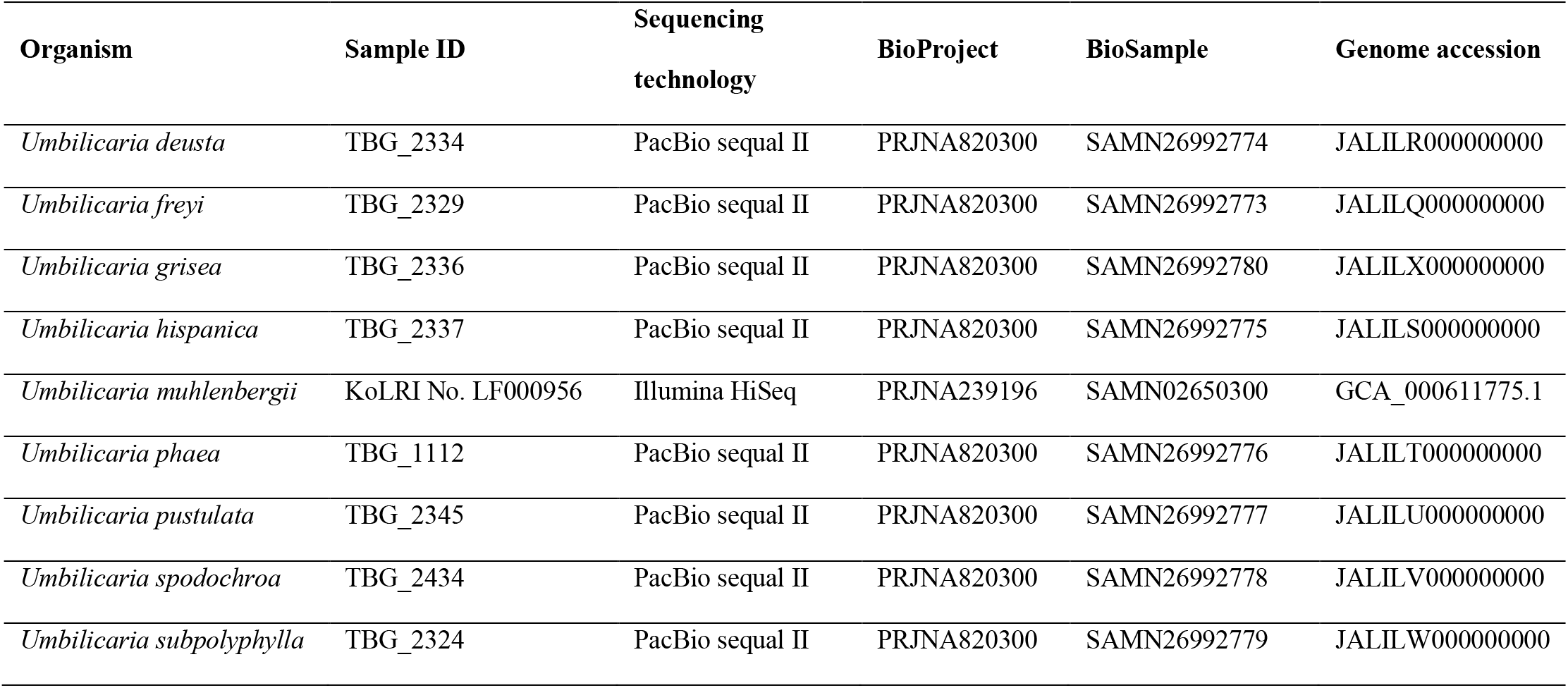
Voucher information of the genomes used in the study.

### BGC prediction and clustering: AntiSMASH

BGCs were predicted using antiSMASH (antibiotics & SM Analysis Shell, v6.0) with scripts implemented in the funannotate pipeline [18,59]. We tested, if a smaller genome size was correlated with a lower number of BGCs. A correlation coefficient near 0 indicates no correlation whereas a coefficient near 1 indicates a positive correlation.

### BGC clustering into BiG-FAM GCFs

The homologous BGCs present in the *Umbilicaria* genomes were grouped into Gene Cluster Families (GCFs) using BiG-FAM, which clusters structurally and functionally related BGCs into GCFs and identifies the structurally most diverse BGCs by comparing the query BGCs to the 1,225,071 BGCs of the BiG-FAM database. The 1,225,071 BGCs in BiG-FAM are clustered into 29,955 GCFs based on similar domain architectures. A GCF comprises closely related BGCs, potentially encoding the same or very similar compounds. By enabling such clustering BiG-FAM establishes the degree of similarity of BGCs of a query taxon to currently known (functionally pre-characterized) fungal and bacterial BGCs. The antiSMASH job ID of each *Umbilicaria* species was used as input for BiG-FAM analysis.

### Quantification of BGC diversity and species specific BGCs in *Umbilicaria*: BiG-SLiCE

We used BiG-SLiCE [20] to identify the most unique or species-specific BGCs within *Umbilicaria*. While BiG-FAM identifies the most diverse BGCs compared to pre-characterized BGCs from other taxa deposited in public repositories, BiG-SLiCE 1.1.0. is a networking-based tool which assesses relations of BGCs of the dataset (i.e., *Umbilicaria* BGCs in our study) and estimates their distance within the dataset to identity unique, species-specific BGCs. The resulting distance indicates how closely a given BGC is related to other BGCs. BiG-SLiCE was run on the *Umbilicaria* BGC dataset (i.e., 217 BGCs from nine *Umbilicaria* spp.) using three different thresholds: 400, 900 and 1800.

## Declarations

### Ethics approval and consent to participate

Not applicable

### Consent for publication

Not applicable

### Availability of data and materials

The dataset(s) supporting the conclusions of this article are available in the GenBank repository, accession PRJNA820300, under the accession numbers JALILQ000000000 - JALILY000000000. The lichen samples of the corresponding *Umbilicaria* Spp. are available as Biosamples SAMN27294873 - SAMN27294881 and the mycobiont samples as Biosamples SAMN26992773 - SAMN26992781. The antiSMASH files of *Umbilicaria* spp. is available at figshare (doi: 10.6084/m9.figshare.19625997).

### Competing interests

None

### Funding

**This research was funded by LOEWE-Centre TBG, funded by the Hessen State Ministry of Higher Education, Research and the Arts (HMWK)**.

### Authors’ contributions

GS analyzed and interpreted the data, generated the figures and tables and wrote the manuscript.

FDG analyzed the data and assisted with the bioinformatic parts of the study.

IS interpreted the data, co-prepared the figures and co-wrote the manuscript.

All authors read and approved the final manuscript.

## Acknowledgements

We thank Prof. Marnix Medema and Dr. Satria Kautsar for their support with BiG-SLiCE program.

## Supporting Information

**S1** Most divergent BGCs in *Umbilicaria* as identified by BiG-FAM, along with the cluster information and sequence.

**S2** Most distantly related BGCs within *Umbilicaria* as identified by BiG-SLiCE along with the cluster information.

## Notes

### Competing Interest Statement

The authors have declared no competing interest.

